# Counting fluorescent emitters with a single photon avalanche diode array

**DOI:** 10.64898/2026.05.01.722215

**Authors:** Clayton Seitz, Carmella Evans-Molina, Jing Liu

**Affiliations:** Department of Physics, Indiana University, Indianapolis, IN 46202, USA; Department of Pediatrics, Indiana University School of Medicine, Indianapolis, IN 46202, USA; Center for Diabetes and Metabolic Diseases, Indiana University School of Medicine, Indianapolis, IN 46202, USA; Herman B Wells Center for Pediatric Research, Indiana University School of Medicine, Indianapolis, IN 46202, USA; Department of Medicine, Indiana University School of Medicine, Indianapolis, IN 46202, USA; Department of Physics and Astronomy, Purdue University, West Lafayette, IN 47907, USA; Purdue Quantum Science and Engineering Institute, Purdue University, West Lafayette, IN 47906, USA

## Abstract

For decades, the photon counting histogram (PCH) was used as the sole method to quantify fluorophore numbers in a diffraction-limited focal volume. This technique combines spatial excitation profiles, and the distribution of photon counts to register the photon emission statistics of individual fluorophores. However, this approach has not yet been transferred to widefield fluorescent imaging due to the lack of fast and single photon sensitive camera sensors which can capture the photon emission statistics of a single fluorophore. Here, we explore avenues towards quantitative analysis of the active fluorophore number by leveraging recent advancements in single photon avalanche diode (SPAD) array technology. Binary exposures of a SPAD array can be synchronized with picosecond laser pulses to measure the PCH in a widefield setting. Then, by modeling the statistical relationship between the active fluorophore number and the PCH in a region of interest following a laser pulse, we can perform Bayesian inference of this number. The model is demonstrated experimentally by counting quantum dots and various numbers of fluorescent dye molecules bound to DNA origamis. We find that this method has several important applications in widefield microscopy, including enhanced localization microscopy and constrained fitting of multiple unresolvable fluorescent emitters.

## INTRODUCTION

Single photon avalanche diodes (SPADs) have long been the detector of choice for highly sensitive biological imaging and sensing, such as single molecule imaging and single molecule spectroscopy. Its fast temporal response and low background noise allow for the registration of a single photon with a temporal resolution as precise as several picoseconds^1^. However, standard SPAD detectors lack spatial resolution and are typically integrated in laser scanning microscopes for biological imaging, which significantly limits the imaging speed. Recently developed SPAD arrays removed this limitation and extended their application to widefield microscopy. SPAD arrays can achieve orders of magnitude higher temporal resolutions than standard cameras, while preserving the SPAD detector’s original signatures, such as single photon sensitivity, low dark count rates, and time-gated photon collection^2–7^. Furthermore, the zero readout noise and large fill-factor of recently commercialized SPAD arrays suggests their use for precision widefield bioimaging for single molecule studies^7^. Such photon number resolving array detectors have also attracted considerable attention in the bioimaging community for applications in fluorescence lifetime imaging^2–4^.

The spatial configuration of the SPAD array can also function as a two-dimensional and “on-chip” Hanbury-Brown and Twiss setup that measures the antibunching effect, which is a purely quantum optical phenomenon that reports the single photon emission signature of a quantum emitter^10^. A simplified SPAD array has been previously discussed for localization microscopy in non-sparse scenes to image quantum dots beyond the diffraction limit^11^. However, this imaging modality is challenged by a general issue encountered in fluorescent labeling, i.e., how to quantify multiple fluorophores in a diffraction-limited spot?

Multiple approaches towards quantification of fluorophores within a diffraction limited spot using SPADs have been proposed in recent years. A classical method based on the confocal setup combined the signature of diffusing dynamics and photon statistics to resolve the distribution of photon numbers collected from a diffraction-limited volume. This photon counting histogram (PCH)^12,13^ analysis yields the number of fluorophores in a small focal volume and provides the quantitative measurement of the diffusion dynamics. Recent utilization of the SPAD array detector in confocal-based fluorescence fluctuation spectroscopy and PCH analysis has significantly extended the feasibility of this classical method towards cutting-edge biological challenges^8,9,14,15^. In addition to resolving the molecule numbers in a dynamic and diffusing environment, the PCH approach was also extended to the application in immobile samples. For instance, the counting by photon statistics (CoPS) approach, counts the number of fluorophores in a diffraction limited volume by combining time correlated single photon counting (TCSPC) with statistical models of the number of multiple detection events observed in the photon counting process^16–18^. Unfortunately, previous modeling approaches for direct fluorophore quantification were designed for beam-splitting confocal configurations and do not directly translate to the SPAD array, where photons can be collected in a time-gated fashion and the probability of a photon arriving at a particular detector element depends on emitter locations^19^.

Here, we propose measurement of the PCH over a region of interest (ROI) for quantitative widefield fluorescent imaging using a SPAD array. In the following sections, we demonstrate that a model of the PCH can be parameterized by the number of active fluorescent emitters in a ROI and their associated molecular brightness. Then, by integrating the PCH into a Bayesian inference scheme, the relative probability of various numbers of fluorescent emitters can be compared, while expressing uncertainty in the molecular brightness apriori. We experimentally demonstrate the model by counting quantum dots and varying numbers of fluorescent dye molecules bound to DNA origamis. In parallel, we use this model to theoretically examine the second-order coherence for various combinations of the expected number of signal photon counts and the expected value of a Poissonian background signal.

## RESULTS

### 1 THE MODEL

#### 1.1 PHOTON STATISTICS OF A FLUOROPHORE UNDER PULSED LASER EXCITATION

Consider the case of laser excitation of an isolated fluorescent emitter in the object plane labeled by a continuous-valued cartesian coordinate θ = (θ_u_,θ_v_) The spatial profile of the intensity in image space due to diffraction is presumed to have a Gaussian shape^20–22^ (Figure 1a).

**Figure 1.**
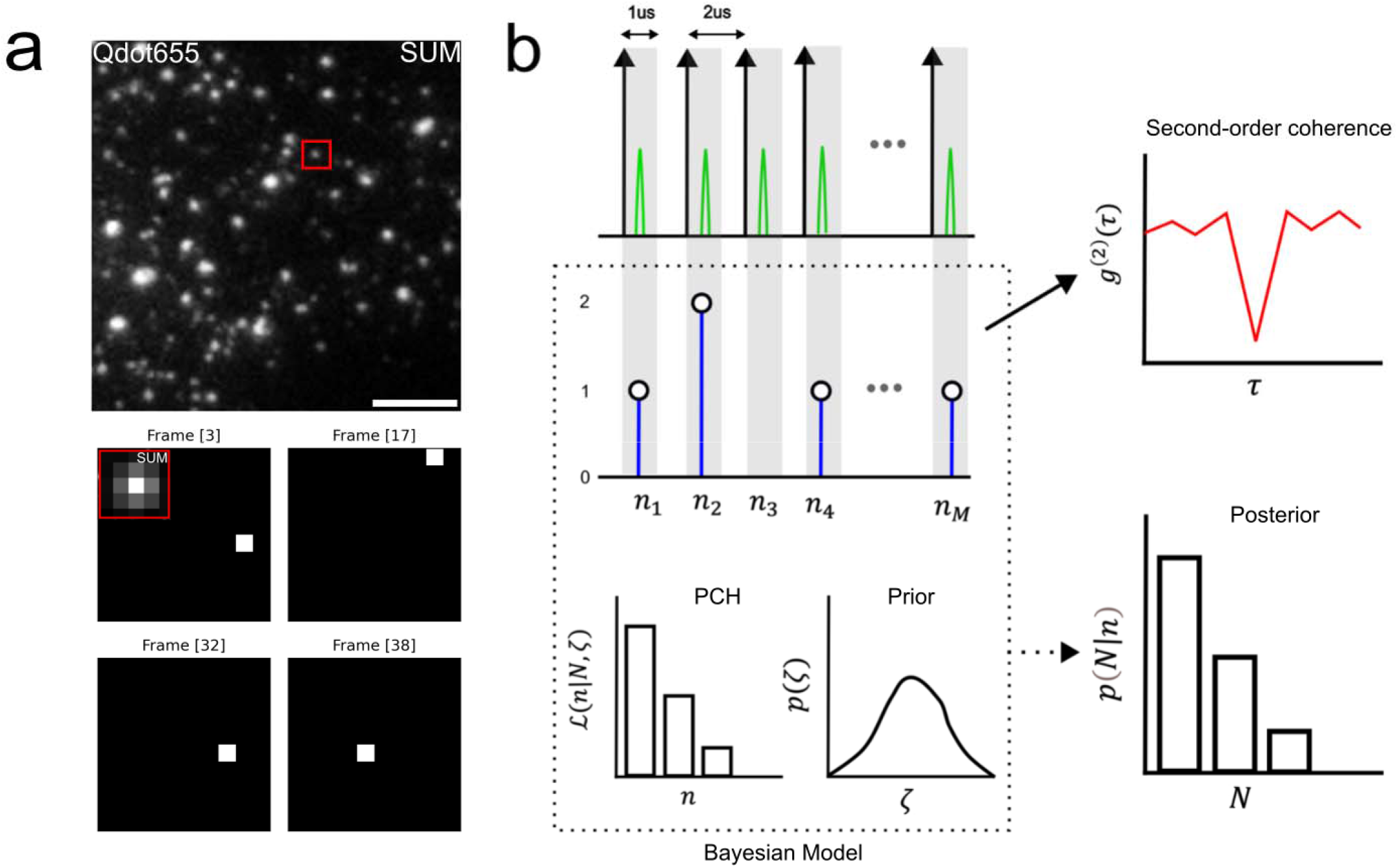
Widefield photon counting and analysis scheme. (a) Image of total photon counts collected from Qdot655 during 30k 1us exposures of the SPAD512 array (b) Single photon imaging scheme using 1us exposures triggered at 500KHz, synchronized with a picosecond laser pulse. The photon counting histogram (PCH) is constructed from the integrated number of photon counts in a region of interest (ROI). Time series of photon counts are used for second-order coherence analysis and the PCH is used in a Bayesian inference scheme of the fluorophore number *N*

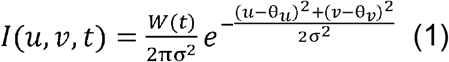

In general, fluorescent emission is a doubly stochastic process^12^, due to temporal fluctuations in the intensity *W*(*t*) as well as the quantum nature of light (Figure 1b). We define the molecular brightness *ζ* (*t*) ∝ *W*(*t*) as the probability a photon is detected from a fluorescent emitter at a time *t*. All factors that affect the photon count rate, such as laser power, absorption cross section, fluorescence quantum yield, detector efficiency, etc., are absorbed into *ζ*(*t*). Furthermore, time-dependence of *ζ* and thus the PCH can arise due to several factors, such as the finite fluorescence lifetime of a fluorophore. Pulsed laser experiments often employ an excitation scheme which removes this source of temporal variability. This is achieved by the condition *τ*_pulse_ ≪ *τ*_fl_ ≪ *τ*_rep_, where *τ*_pulse_ is the duration of the laser pulse, *τ*_fl_ is the fluorescence lifetime, *τ*_rep_ and is the laser repetition time^16^. Other sources of temporal variability lead to a distribution *π* (ζ) in the molecular brightness such that ζ(*t*) ∼*π*(ζ). In this case, the probability of detecting a photon from a fluorophore after a laser pulse simply becomes the mean molecular brightness 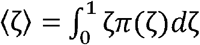 and the PCH remains stationary. From here on, we refer to the time-independent mean molecular brightness as ζ or simply as molecular brightness. Additional justification for this treatment is given in the Supplement.

Suppose we use this scheme to measure the PCH over a ROI bounding all signal photon counts for an isolated single photon source. In this scenario, the spatial variation of the intensity given in Eq. (1) can be ignored, and the PCH will follow a Bernoulli distribution:

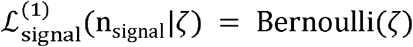

More generally, for *N* fluorophores emitting photons which can be detected within a ROI of the SPAD array, the number of signal photons measured n_signal_ following a single excitation pulse will have Binomial statistics n_signal_∼Binom(*N*,ζ). The fluorophore number *N* enters here as the number of trials run during a single excitation pulse. For a Poissonian background signal, the number of background counts reads n_bg_ ∼ Poisson (*λ*) for an expected number *λ* of background counts in the ROI per frame. In our experiments, we can measure *λ* directly, as the background signal arises from sensor noise and a small of portion of the excitation laser light. The total number of counts n = n_signal_ + n_bg_ detected in the region of interest following a single pulse is then distributed by the following PCH:

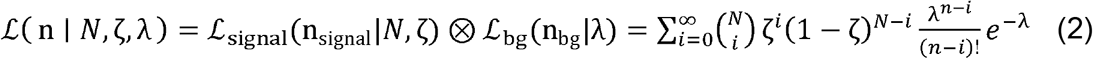

This expression represents a convolution of Poisson and Binomial probability mass functions and is the primary means of inference of the number of active emitters *N* in a ROI as well as theoretical analysis of the second-order coherence.

#### 1.2 SECOND ORDER COHERENCE

A time series of fluorescence photon counts measured by a SPAD can have a temporal correlation structure, which is generally quantified by the second-order coherence function. However, for pulsed excitation schemes where *τ*_pulse_ ≪ *τ*_fl_ ≪_rep_, the temporal correlation structure of fluorescence photons due to the fluorescence lifetime is eliminated. As a result, the number of photon counts measured in a diffraction limited volume following each laser pulse is an independent and identically distributed sample from the stationary PCH, significantly simplifying our theoretical analysis of the second-order coherence.

To measure second-order coherence in experiments, the normalized second order coherence function *g*^(2)^(*m*) is used^11^. An empirical estimate of *g*^(2)^(*m*) is made, based on the number of coincidences *G*^(2)^(*m*) at a lag time *m* in the ROI:

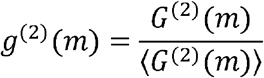

where ⟨*G*^(2)^ (*m*) ⟩ denotes a time average over a signal shifted by lag time *m*. In words, the function *G*^(2)^(*m*) counts the number of coincidences at a lag time *m* and is normalized by its expectation ⟨*G*^(2)^ (*m*) ⟩. It is well known that, for Poissonian light, the second order coherence function *g*^(2)^(*m*) should be approximately unity for all lags *m*. This is not necessarily the case, however, for binomial states^23^ of the quantized radiation field or under all experimental conditions tested here. To investigate the properties of the *g*^(2)^ (0) dip, we use the PCH to identify distinct distributions for *G*^(2)^ (0) and *G*^(2)^ (*m*) for nonzero lags *m* (see Methods). For example, we find that the PCH ℒ(n|*N*,ζ,*λ*) parameterizes a probability distribution on *G*^(2)^ (0) and therefore *g*^(2)^ (0). These distributions can be then used to numerically estimate ⟨*g*^(2)^ (0)⟩ and its variance under different conditions. Importantly, we found significant heterogeneity in the value of ⟨*g*^(2)^ (0) ⟩ as the molecular brightness and background rate covary. Indeed, high signal to background ratios give the traditional *g*^(2)^(0) dip expected for a binomial signal; however, ⟨*g*^(2)^ (0)⟩ can be below unity even in the absence of signal i.e., pure Poissonian photon statistics (Figure 2a). This can be seen from that for pure background signal, low background rates give few coincidences, similar to a single photon source. In our analysis, we find that larger background rates under signal-free conditions give a ⟨*g*^(2)^ (0)⟩ of unity. Furthermore, ⟨*g*^(2)^ (0) ⟩ can saturate for larger values of the number of emitters for larger combinations of the molecular brightness and background rate (Figure 2b).

**Figure 2.**
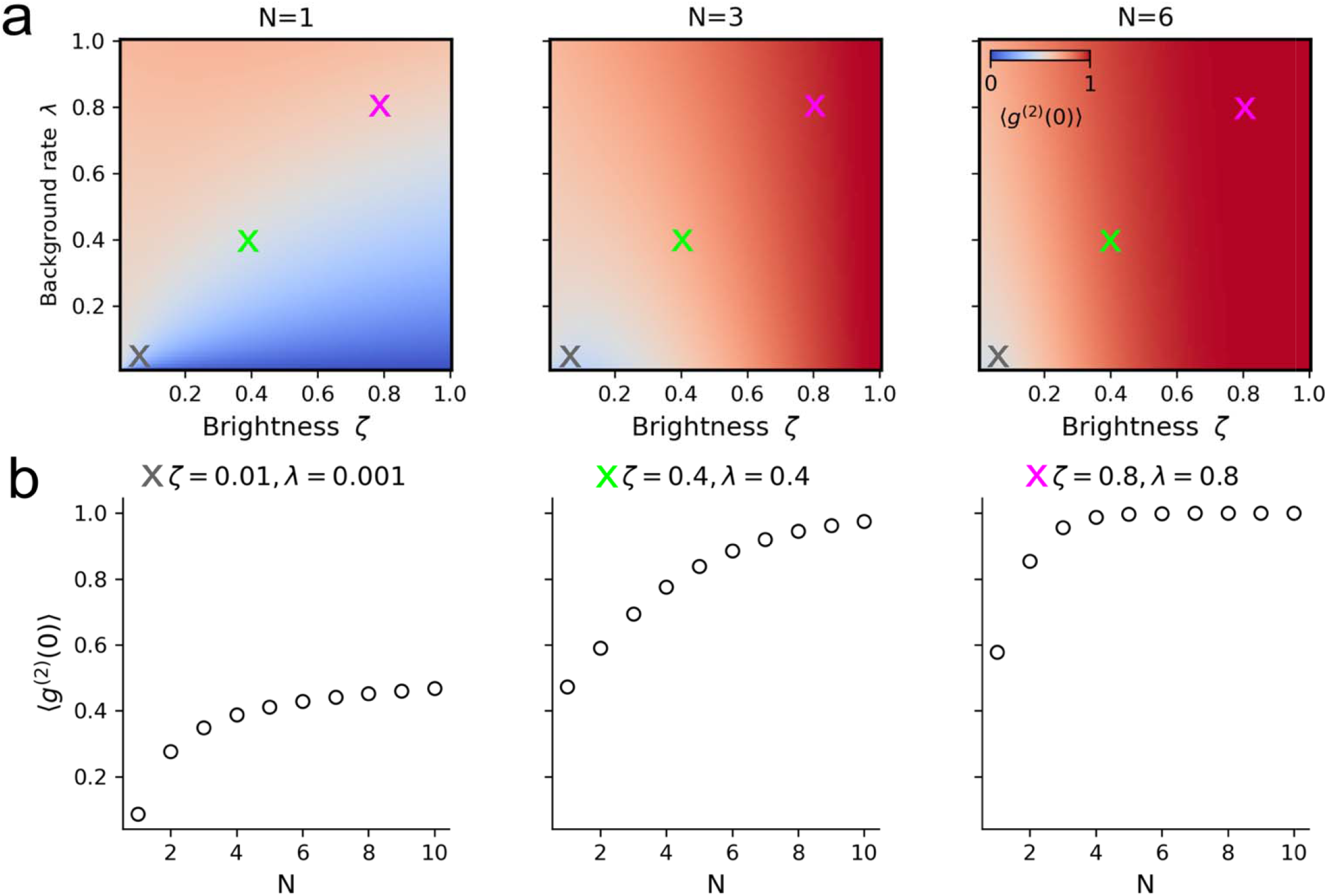
Second-order coherence behavior under the PCH model. (a) Heatmap of the expected value of for various combinations of the molecular brightness (photon detection probability) and background rate parameter. (b) Expected value of for selected combinations of the molecular brightness and background rate parameter as a function of the number of fluorescent emitters.

#### 1.3 BAYESIAN INFERENCE OF THE NUMBER OF ACTIVE EMITTERS

When the number of fluorescent emitters *N* is unknown, the PCH ℒ(n|*N*,ζ,*λ*) can be used in a Bayesian inference scheme to construct a posterior distribution on the binomial parameters:

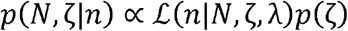

We use a Gaussian prior on ζ i.e., 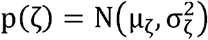 and presume the background rate *λ* is a known parameter based on experimental measurement. The joint posterior can be integrated ζ over to produce a posterior distribution on the fluorophore number *N* i.e., 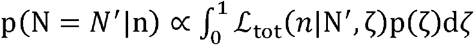 with 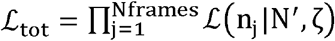. This total likelihood is made tractable by a log-sum exponential trick 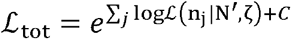 for a constant *C* determined empirically. The posterior distribution *p*(*N*|*n*) is obtained by Monte Carlo integration of the joint posterior *p*(*N*,ζ|*n*) over ζ. This involves sampling many ζ values from the prior and, for each sampled ζ the PCH is computed. These PCH values are then weighted by the prior probabilities *p*(ζ). The result is obtained by averaging these weighted likelihoods over all sampled ζ, which approximates the integral:

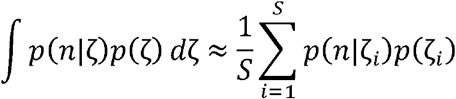

where *S* is the number of samples from the prior. The final posterior is then estimated by minibatching the data into batches of 1000 frames and averaging the posterior p(N|n) over minibatches. Estimates of the fluorophore number *N* within each ROI can be found from the maximum aposteriori (MAP) estimate N* given by p(N|n).

### 2. MODEL VALIDATION WITH SIMULATION DATA

To validate our model, we apply it to simulated photon emissions from single and multiple quantum emitters. Simulated photon count time-series were generating by sampling from the PCH using various values for *N* and setting *ζ* =0.01 (Figure 3a,e,i). Photon counting histograms show an increased probability of photon coincidence with increasing values of the fluorophore number (Figure 3b,f,j). As expected, we found that *g*^(2)^ (0) < 0.5 while for increasing values of the fluorophore number *g*^(2)^ (0) approached *g*^(2)^ (0) = 0.5 (Figure 3c,g,k). Analysis of the posterior distribution on the fluorophore number successfully recovered the value of the fluorophore number used to parameterize simulations (Figure 3d,h,l). Moreover, increasing values of the fluorophore number results in a posterior distribution with decreasing precision, indicating a loss of certainty in the true fluorophore number. Therefore, we find that MAP estimates of the fluorophore number can be error prone for higher values of the fluorophore number. A complete analysis of the error and bias in MAP-based fluorophore number estimation is shown in (Supp. Figure 1).

**Figure 3.**
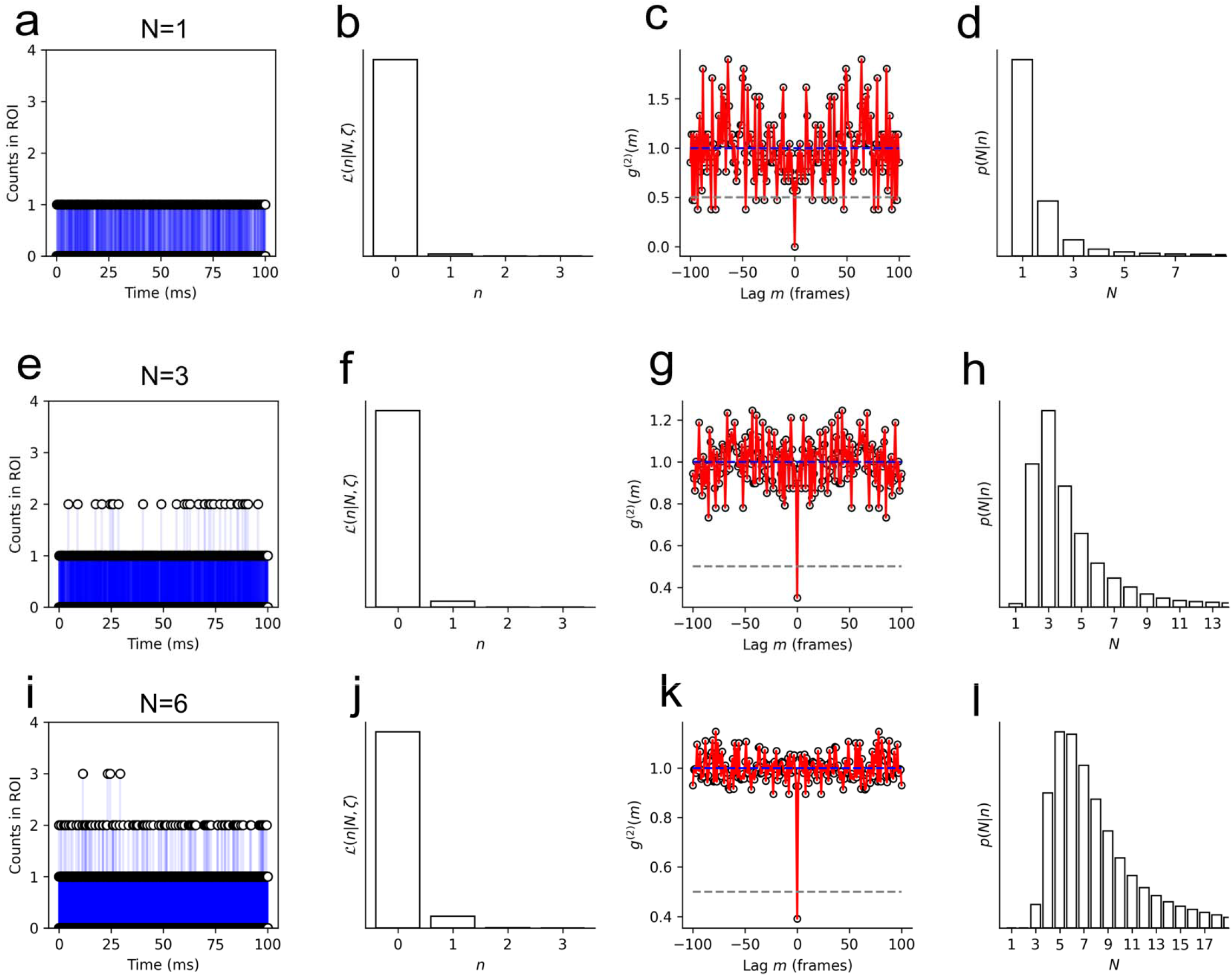
Photon counting histogram and analysis on zero-background simulations (a) Simulated photon counts in 1us exposure for N=1 and (b) Photon counting histogram for counts in (a) (c) Second-order coherence function for data in (a) (d) Posterior distribution for data in (a). (e) Simulated photon counts in 1us exposure for N=1 and (f) Photon counting histogram for counts in (e) (g) Second-order coherence function for data in (e) (h) Posterior distribution for data in (e). (i) Simulated photon counts in 1us exposure for N=1 and (j) Photon counting histogram for counts in (i) (k) Second-order coherence function for data in (i) (l) Posterior distribution for data in (i). Inference in (d,h,l) use

### 3. DISTINGUISHING SINGLE QUANTUM DOTS FROM ASSEMBLIES

A custom widefield fluorescence microscope was built for widefield photon counting by synchronization of laser pulses with 1-bit exposures of a SPAD array. This excitation scheme permits the computation of the second order coherence function *g*^(2)^(*m*) and visualization of the spatial intensity distribution by summing photon counts over time. The acquisition scheme and analytical methods were then applied in two experimental contexts. First, we examined the posterior distribution and *g*^(2)^(*m*) function for putatively isolated quantum dots as well as quantum dot aggregates. Quantum dots exhibit temporally heterogeneous photoluminescence (PL) due to nonradiative transitions of electrons in the conduction band giving rise to a phenomenon known as blinking^24,25^ Quantum dots showing clear two-state PL fluctuations exhibited *g*^(2)^ (0) < 0.5 and a posterior distribution maximized at *N* = 1 (Figure 4a-d). Aggregates of quantum dots with more complex PL fluctuations showed *g*^(2)^ (0) = 0.5 and a posterior distributed around larger values of the fluorophore number (Figure 4e-h).

**Figure 4.**
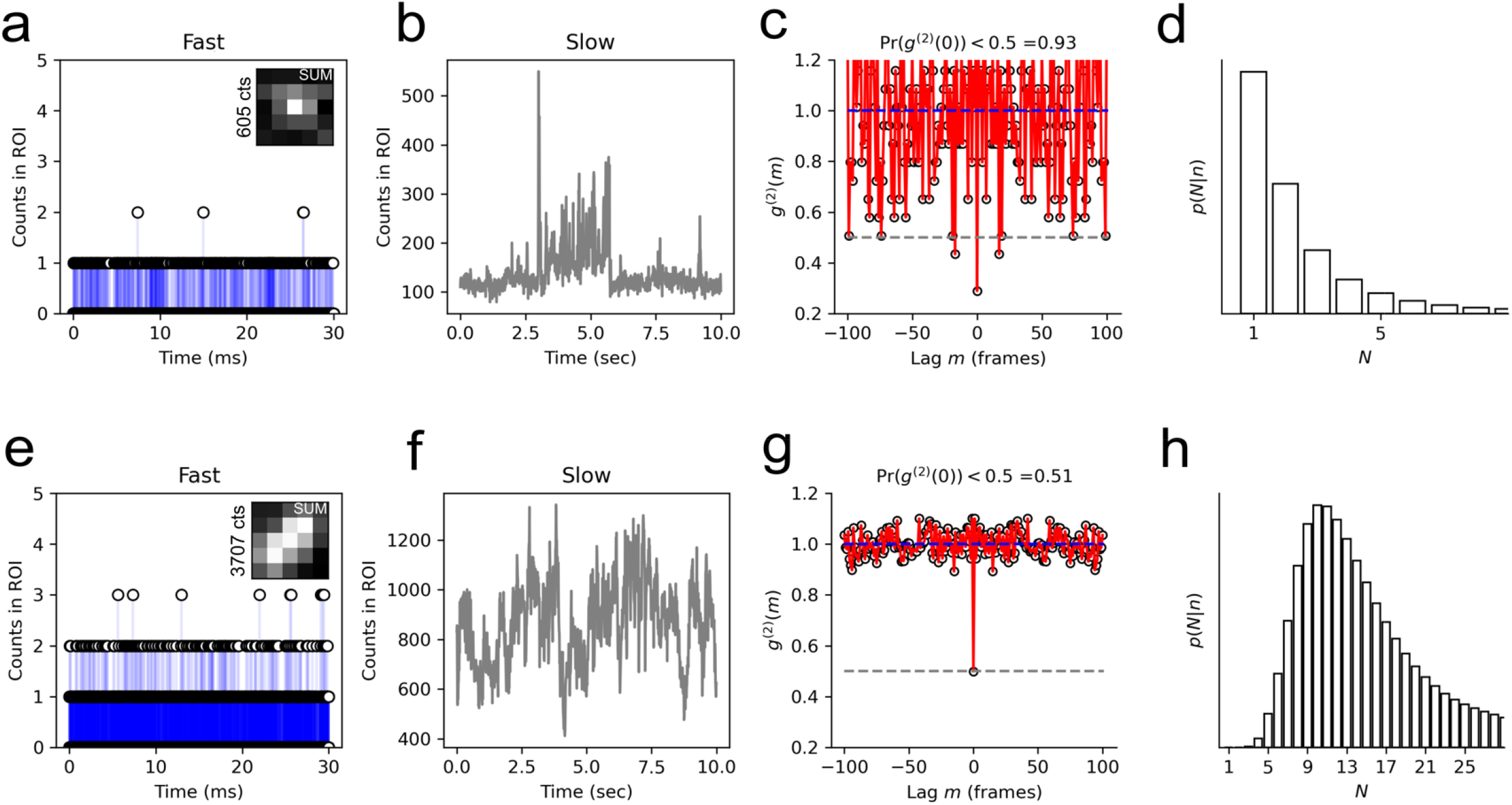
Distinguishing single and multiple quantum dots (a) Photon counts in 1us exposure using 532nm pulsed excitation of a putatively isolated QD (b) Photon counts in 10ms exposure using 488nm continuous-wave excitation (c) Second-order coherence function for data in (a) (d) Posterior distribution for data in (a) (e) Photon counts in 1us exposure using 532nm pulsed excitation of a QD aggregate (f) Photon counts in 10ms exposure using 488nm continuous-wave excitation (g) Second-order coherence function for data in (e) (h) Posterior distribution for data in (e). Inference parameters can be found in Supplementary Table 1.

### 4. COUNTING FLUOROPHORES BOUND TO DNA ORIGAMIS

In a second series of experiments, we examined the posterior distribution and *g*^(2)^(*m*) function for DNA origamis which can bind up to *N* = 3 or *N* = 6 ATTO532 fluorescent dye molecules, referred to as G3 and G6, respectively (Figure 5a,e). For origamis which can bind up to *N* = 3 ATTO532 dyes, no clear *g*^(2)^ (0) dip was observed, due to the low signal to background ratio at this signal level (Figure 5c). For origamis which can bind up to *N* = 6 ATTO532 dyes, no clear *g*^(2)^(0) dip was observed, due to the large *N* character of the *g*^(2)^ (0) dip predicted by the theory (Figure 5g). We conclude that the *g*^(2)^ (0) dip may lead to ambiguous interpretations at low signal levels. However, the posterior distribution can estimate the *N* value from the photon count distribution for G3 (Figure 5d) and G6 (Figure 5h). Indeed, distributions of the maximum aposteriori (MAP) estimate of the fluorophore number showed consistency with the known value of *N* for G3 and G6 (Figure 6).

**Figure 5.**
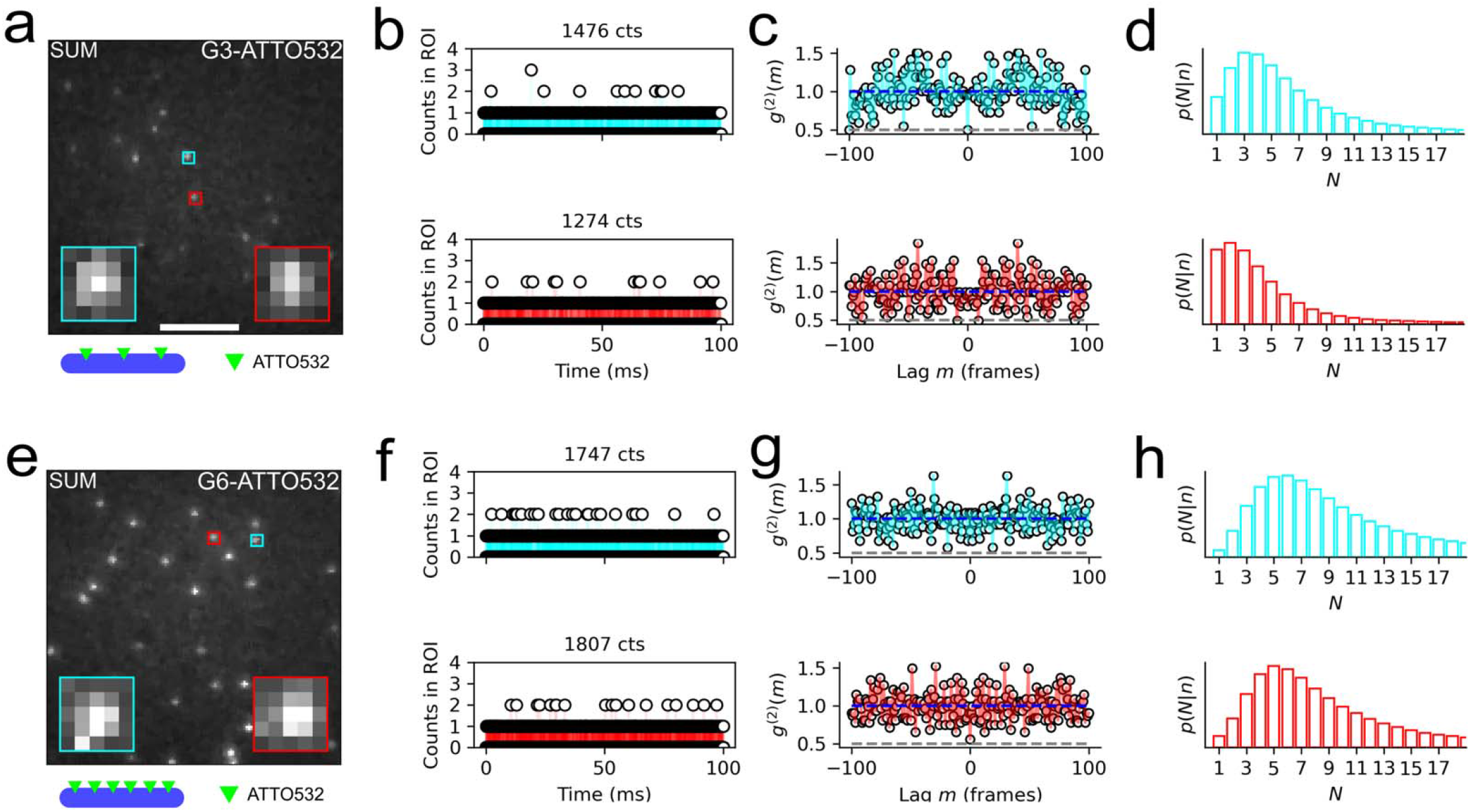
Counting ATTO532 dye bound to DNA origamis. (a) Widefield image of summed photon counts collected from DNA origami with three ATTO532 binding sites (G3 sample). (b) Photon counts in 1us exposures using 532nm pulsed excitation of G3 sample for two example spots (c) Second order coherence for two example spots in G3 (d) Posterior distribution for two example spots in G3 (e) Widefield image of summed photon counts collected from DNA origami with three ATTO532 binding sites (G6 sample) (f) Photon counts in 1us exposures using 532nm pulsed excitation of G6 sample for two example spots (g) Second order coherence for two example spots in G6 (h) Posterior distribution for two example spots in G6. Inference parameters for G3 and G6 can be found in Supplementary Table 1.

**Figure 6.**
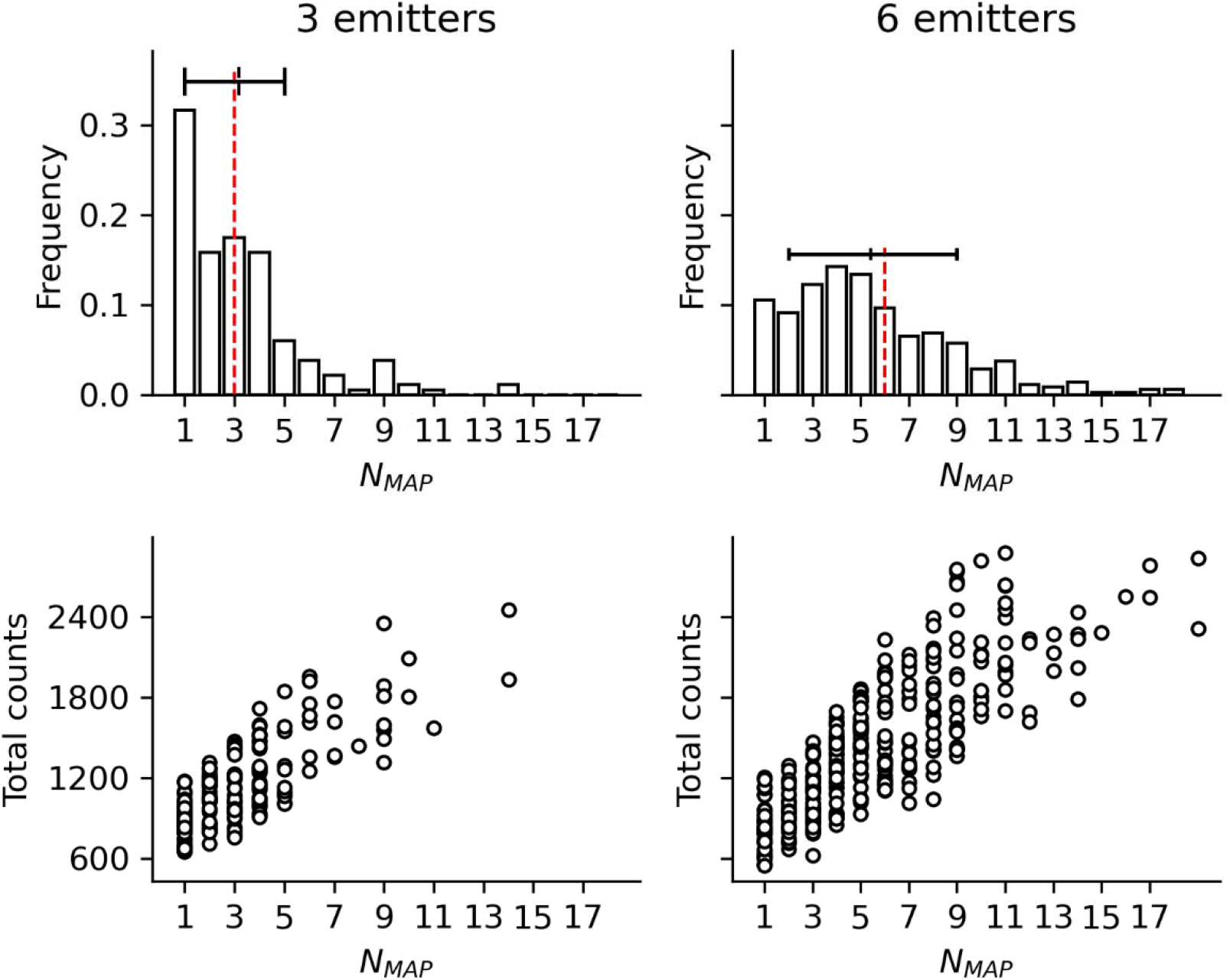
Distributions of the maximum aposteriori estimate of the number of ATTO532 dye bound to DNA origamis (left) Distribution of estimate of the fluorophore number and total photon counts for DNA origamis with three ATTO532 binding sites (G3 sample, 194 spots used for analysis) (right) Distribution of estimate of the fluorophore number and total photon counts for DNA origamis with six ATTO532 binding sites (G6 sample, 372 spots used for analysis). Above each histogram a box plot indicates the central 68% quantile about the median of the probability density. The red dashed line represents the expected emitter number.

## DISCUSSION

In this paper, we leveraged the recently developed SPAD array to spatially resolve the photon counting histogram with a widefield microscope. The PCH was integrated into a Bayesian inference scheme to extract the number of fluorescent emitters throughout the field of view and measure the second order coherence function. We demonstrated counting of fluorescent quantum dots and fluorescent dyes bound to DNA origami, suggesting that this is a capable method for quantitative widefield fluorescent microscopy. Future work may assess more complex fluorescent imaging scenarios.

For example, fluorescence intermittency or photobleaching leads to a *ζ*(*t*) which is not constant and can be fluorophore specific. This represents another source of temporal variability in the observed photon-number fluctuations and the PCH function ℒ_signal_ (n_signal_). Therefore, due to the unique dynamics of *ζ*(*t*) for each fluorophore, the use of equation (2) is no longer appropriate, and one can expect more complex behavior of *g*^(2)^ (0). Similarly, challenges may arise in distinguishing one homogeneously emitting fluorophore and several blinking fluorophores. If the effect of censoring photons by blinking can be accounted for, the technique used here may be compatible with common nanoscopy methods which rely on fluorescence intermittency. However, neither of these effects need to be considered here. The tested quantum dots are photostable and photobleaching of ATTO532 could be mitigated under the experimental conditions used. The on and off state lifetime of tested quantum dots and are significantly longer than the acquisition sequence, making fluorescence intermittency negligible.

The acquisition times necessary to obtain sufficient photon counts for computing the necessary statistics can potentially be short. Most fluorophores have relaxation times in the nanosecond range and thus photons can be collected at a rate of tens of millions of excitation pulses per second. These rates are currently difficult to obtain, however, due to limitations in detector throughput. Current exposure times of the SPAD array are limited to hundreds of nanoseconds to microseconds, precluding detailed temporal analysis of the second-order coherence. Moreover, the data volume may become intractable due to the need for several thousands of frames for a millisecond-scale exposure time.

Meanwhile, the effect of optical crosstalk between two SPAD array elements would lead to some “false-positive” results for the photon statistics^26-28^. This can occur when an avalanche is triggered in the “emitter” followed by emission of a second photon, which can trigger an avalanche in another SPAD, referred to as the “detector”. We have derived the optical crosstalk probability of the SPAD array detector (Supplementary Materials), and have experimentally measured that with a classical light source. Similar as the result reported by other groups^29^, we have observed negligible crosstalk probability (<4.6×10^-4^) between neighboring pixels (Supp. Figure 2) and such crosstalk was undetectable in our photon counting experiments.

Theoretical analysis of the *g*^(2)^ (0) value under the PCH model demonstrates that the measured value can strongly depend on experimental conditions. For example, error in the *g*^(2)^ (0) is dependent on the number of photons collected (Supp. Figure 1). The sensitivity *g*^(2)^ (0) complicates its direct interpretation in quantitative widefield microscopy; however, *g*^(2)^ (*m*) may reveal characteristics of the temporal autocorrelation of photon counts. Sensitivity of *g*^(2)^ (0) to experimental conditions ultimately requires a statistical model of *g*^(2)^ (0), which under the pulsed excitation scheme used here, is the PCH. Therefore, the statistical inference scheme presented here allows us to account for this dependence and perform quantitative widefield microscopy over a range of experimental conditions.

Lastly, the method proposed here may lead to significant advances in localization microscopy. Many of these schemes utilize the concept of precise localization of fluorescent emitters to produce super-resolved images^30,31^. However, an inherent problem with such methods is the assumption that fluorescent emitters are isolated, which can lead to undercounting and localization errors. Various models for multi-emitter localization have been developed to approach this issue statistically^32–36^, which necessarily treat the number of fluorescent emitters as an unknown. The approach proposed here provides a physical means to quantifying active fluorescent emitters based on widefield photon counting.

In conclusion, we propose a single molecule imaging technique that allows for counting of fluorescent molecules by modeling the quantum properties of fluorescence emission. The technique does not require a nonclassical light source and is designed to supplement standard widefield microscopy techniques towards quantitative fluorescence imaging.

## ACKNOWLEDGEMENTS

Authors would like to acknowledge the funding support from the National Institute of General Medical Sciences (NIH 1R35GM147412), National Institute of Diabetes and Digestive and Kidney Diseases (NIH 1R03DK135457 to JL; R01 DK127308 to CEM). This project has been made possible in part by grant number 2023-328664 from the Chan Zuckerberg Initiative DAF, an advised fund of Silicon Valley Community Foundation.

## MATERIALS AND METHODS

### QUANTUM DOTS AND DNA ORIGAMIS PREPARATION

Fluorescent samples used here were either Quantum dots (Qdot655, ThermoFisher) coated on a glass coverslip, or ATTO532 tagged DNA origamis (GATTAquant). Fluorophores in a region with quasi-uniform laser power were selected for analysis to simplify the prior distribution on the molecular brightness. DNA origami samples are a commercially made sample containing origamis with either three or six ATTO532 binding sites for testing the Bayesian inference scheme and second-order coherence analysis.

### SIMULATION OF PHOTON EMISSION RECORDED BY THE SPAD ARRAY

Simulated photon count time-series were generating by sampling from the PCH in equation (2) with the Python programming language using variable values for N and *ζ*=0.01. The second-order coherence *g*^(2)^(*m*) and posterior distributions were computed as described previously.

### SINGLE MOLECULE IMAGING WITH THE SPAD ARRAY

A 512×512 SPAD array (Pi Imaging Technologies) was connected to the custom-built single molecule imaging system based on the ASI RAMM. A picosecond 532nm pulsed laser (Picoquant) was triggered at 500 kHz as the excitation source. A complete schematic of the imaging system is given in (Supp. Figure 3) Laser power was set at 300uW at the back focal plane of the microscope objective for all experiments. Emission light was collected using an oil-immersion 100X/1.4NA objective (Nikon). The emission signal was filtered to exclude the laser line (Semrock) and projected onto the sensor with the tube lens. The acquisition of the SPAD array is synchronized with the pulsed laser frequency. Each acquisition consists of a series of 1-bit frames, using a 1us exposure per frame. For quantum dot imaging, 30ms total exposure was used (30k frames). For DNA origami imaging, 100ms total exposure was used (100k frames). To obtain time-course data of photon counts, we (i) summed binary images over the entire acquisition (ii) estimated spot centroids from summed binary images using the Laplacian of Gaussian (LoG) detection algorithm, and (iii) extracted time-series of photon counts in a 5×5 pixel region of interest around each detected spot.

### COMPUTATION OF THE SECOND ORDER COHERENCE

In practice, we compute the second order coherence at zero lag using

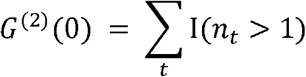

where I is an indicator function. The value of this function at nonzero lag is given by

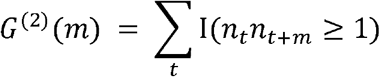

The normalized second order coherence *g*^(2)^ (*m*) is then computed over a range of lags − *m*_*min*_ ≤ *m* ≤ *m*_*max*_. Theoretical estimates of *g*^(2)^ (0) can be obtained by determining the generating model of *G*^(2)^ (*m*). We find that for a sequence of *N*_frames_, the PCH can be used to derive Binomial distributions for *G*^(2)^ (0) and *G*^(2)^ (*m*)

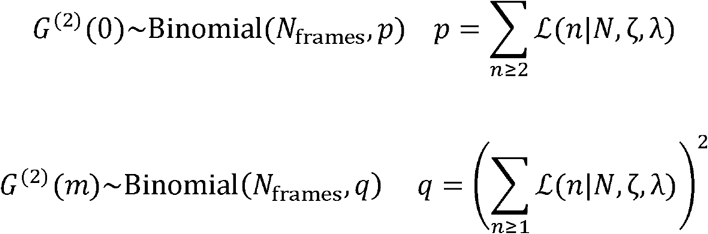

The expected value of *g*^(2)^ (0)can be derived from these distributions:

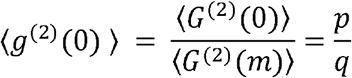

Error in the estimate of *g*^(2)^ (0) is found by the expression

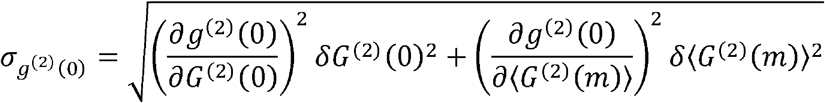

This expression simplifies considerably in the case that *δ* ⟨ *G*^(2)^ (*m*)⟩ = 0 i.e., certainty in ⟨ *G*^(2)^ (*m*)⟩

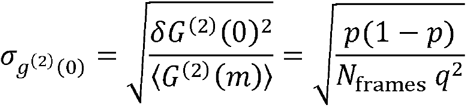

The value of 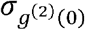 is therefore a function of *ζ* as well as the number of frames in the acquisition *N*_frames_.

